# Proline 110 is necessary for maintaining a compact helical arrangement in caveolin-1

**DOI:** 10.1101/2025.07.10.664188

**Authors:** Katrina Brandmier, Kerney Jebrell Glover

**Author notes:** Correspondence, Kerney Jebrell Glover, Department of Chemistry, Lehigh University, 6 E. Packer Avenue, Bethlehem, PA 18015, USA.

## Abstract

Caveolin-1 (Cav1) is an integral membrane protein essential for the formation of caveolae, plasma microdomains implicated in signal transduction and mechanoprotection. Cav1 is comprised of three major alpha helices, but the topology these helices adopt remains unclear. Proline 110 is located between helix 1 and helix 2, and is hypothesized to enable Cav1 to adopt an intramembrane turn crucial for the cytosolic topology of Cav1. To assess the structural role of Proline 110, we utilized Förster resonance energy transfer (FRET) between native tryptophan (W128) and site-specifically labeled dansyl fluorophores to monitor conformational changes induced by the mutation of Proline 110 to Alanine (P110A). Static light scattering confirmed that all FRET constructs behaved monomerically, ensuring intramolecular energy transfer measurements. Our results show a significant decrease in FRET efficiency upon the P110A mutation consistent with a large conformational change. These findings support the critical role of P110 in maintaining the native topology of Cav1 and highlights the structural sensitivity of the intramembrane turn.

## Introduction

Caveolins are a family of integral membrane proteins that are crucial for the formation of caveolae, flask-shaped invaginations of the plasma membrane. Caveolin has three isoforms (-1,-2,-3) which share structural features but differ in function (1–3). Caveolin-1 (Cav1), the most broadly expressed isoform, is required for the formation of caveolae in non-muscle cells such as adipocytes, endothelial, and fibroblasts (4–9). Beyond its structural role in caveolae, Cav1 has been implicated in diverse cellular processes including endocytosis, signal transduction, and mechano-protection (10–13). Despite being only 178 amino acids, Cav1 adopts a compact, tightly membrane-associated topology that has yet to be fully elucidated.

Structural studies indicate that Cav1 inserts into the cytoplasmic leaflet of the membrane and contains three a-helical regions. Helices 1 and 2 (H1 and H2), which are proposed to insert deeply into the membrane, and Helix 3 (H3), an amphipathic helix that lies parallel to the membrane (14). The N- and C-termini of Cav1 reside on the cytoplasmic side of the membrane, making it a monotopic membrane protein (15). While this structural framework is broadly accepted, the precise membrane topology of Cav1 remains unresolved. This unique topology has led to the hypothesis that Cav1 forms a tight intramembrane turn between H1 and H2, allowing it to anchor into the cytoplasmic leaflet without spanning the bilayer. A similar helical arrangement has been proposed for Cav3 (16). Supporting this model, molecular dynamics (MD) free energy simulations of H1 and H2 have shown that a U-shaped conformation is energetically viable within 1,2-dimyristol-*sn*-glycero-3-phosphocholine (DMPC) bilayer (17).

Within Cav1, H1 and H2 are separated by three residues, GIP, where Proline 110 (P110) is highly conserved and located near the midpoint of the H1-H2 segment. Prolines are known to disrupt α-helices, making them ideal candidates for facilitating sharp turns within the hydrophobic core of membranes. The position of P110 and its conservation suggest it may serve as a hinge that enables the confirmation between H1 and H2 to adopt an intramembrane turn. This hypothesis has been explored by Epand et al. where the structural role of Proline 110, part of the putative intramembrane turn, was explored (18). They found that expression of Cav1 with a FLAG-tag at the N-terminus in HEK293 cells resulted in no antibody activity at the exterior cell surface, consistent with cytosolic localization of the N-terminus. However, the mutation of proline 110 with alanine (P110A) rendered the N-terminus antibody-accessible at the exterior cell surface, suggesting a major topological rearrangement. Computational modeling supported this, showing that the wild-type hydrophobic segment adopts a U-shaped conformation, while the P110A mutation straightens this segment into a single long transmembrane helix (18). However because the location off the C-terminus of Cav1 was not probed in the study, the precise effect of this mutation—whether it causes helix linearization as suggested by the MD simulations or another distinct topology with an extracellular N-terminus —remains experimentally unresolved (Figure 2).

To address this ambiguity, we employed Förster resonance energy transfer (FRET) to monitor conformational changes in Cav1. FRET is a phenomenon where energy can be transferred between a donor fluorophore and an acceptor fluorophore, as long as the excitation of the acceptor fluorophore overlaps spectrally with the emission of the donor fluorophore. FRET is highly distance-dependent and can measure distances between 10 Å and 100 Å (19). Herein, we utilize tryptophan (a native residue) as the donor and dansyl as the acceptor fluorophore (Figure 1). Our FRET analysis revealed a significant loss of energy transfer in the P110A mutant compared to wild-type, indicating a substantial conformational change associated with the mutation and further reinforces the critical role of P110 in maintaining the native topology of Cav1.

**Figure 1.**
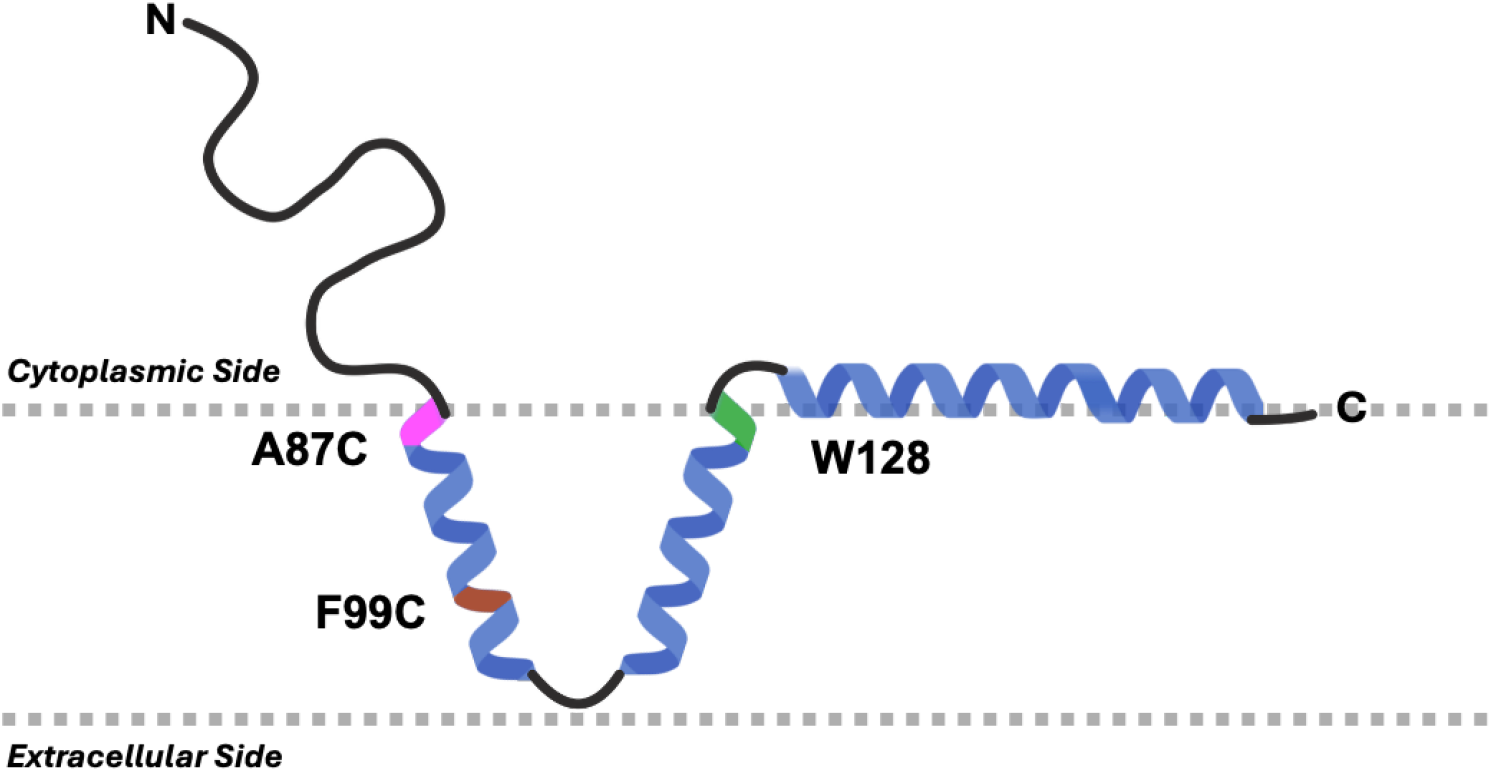
Cartoon hypothesized membrane topology of Cav1, red highlights site-specific acceptor positions, and green highlights donor position.

## Materials and Methods

Empigen BB^®^ was purchased from Sigma-Aldrich (St. Louis, MO). Ni Sepharose 6 Fast Flow resin and Sephacryl^®^ S300 HR 16/60 were purchased from Cytiva Life Sciences (Marlborough, MA). 1,5-IAEDANS was purchased from Setareh Biotech (Eugene, OR). 1-Naphthalenesulfonamide, 5-(dimethylamino)-N-[2-(2-pyridinyldithio)ethyl] (dansyl PDS) was synthesized in-house. Lysozyme was purchased from ThermoFisher Scientific (Waltham, MA). Tris(2-carboxyethyl)phosphine hydrochloride (TCEP) was purchased from GoldBio (St. Louis, MO). 96 well plates were purchased from Greiner Bio-One (Monroe, NC). Amicon^®^ Ultra centrifugal concentrators were purchased from MilliporeSigma (Burlington, MA). All other reagents were of standard ACS grade.

### Caveolin-1 FRET mutagenesis

For FRET experiments tryptophan 128 (W128) is used as the donor position as it rests at the C-terminus of H2, an ideal position for probing the intramembrane turn. To ensure specificity of observed FRET, all other tryptophan residues were conservatively mutated to phenylalanine (W85F, W98F, W115F). The acceptor position was located at residue 87 (A87C), positioned at the N-terminus of H1, or at residue 99 (F99C) located near the center of H1 (Figure 1). Cav1 is natively palmitoylated at three cysteine residues C133, C143, and C156. However, mutations of these cysteines to serines has been shown to have no effect on the trafficking of Cav1 to the plasma membrane. Therefore, to design the FRET constructs, the following point Cav1 constructs were made:

A87C W128

A87C P110A W128

W128

W128 P110A

F99C W128

F99C P110A W128

All mutations were made using the QuikChange^®^ Site-Directed Mutagenesis Kit purchased from Agilent (Santa Clara, CA).

### Caveolin-1 expression and membrane preparation

Cav1 with a C-terminal myc tag was cloned in the pET-24a(+) vector appending a C-terminal hexahistidine tag, and transformed into BL21(DE3) *E. coli* cells for overexpression using the autoinduction method by Studier *et al*. (20) In short, 1000 µL of an overnight (20 h) starter culture in MDG media was used to inoculate 1 L of ZYM-5052 media in a 6 L Erlenmeyer flask. The culture was shaken at 25°C for 24 hours at 250 rpm. Cells were harvested by centrifugation at 8,200 x *g* for 30 min at 4°C in 4 × 250 mL centrifugation bottles. Cells were then washed with 0.9% (w/v) NaCl and recentrifuged. Cell pellets were stored at -20C until ready for use. One 250 mL pellet was resuspended in 25 mL of 1x TAE (40 mM Tris-acetate, 1 mM EDTA, pH 8.0) supplemented with 1 mg/mL lysozyme and stirred at 4°C for 1 hour, then sonicated using a Branson sonifier 450 equipped with a flat tip (duty cycle 3, output power 3) until the cells were fully homogenized (∼15 min). The temperature was kept below 10°C during the sonication process. The homogenized cells were then centrifuged at 7000 x *g* for 30 minutes to pellet cellular debris. Carefully, the supernatant was removed and the membranes pelleted by centrifugation at 230,000 x g for 20 minutes at 4°C. Membranes were then washed by re-homogenizing each membrane in 6 mL of 1× TAE using a 3 mL syringe and 21 gauge x 1 (0.8mm x 25mm) needle. 0.5 mM TCEP was present for all cysteine-containing constructs. Washed membranes were re-pelleted by centrifugation at 230,000 x g for 20 minutes at 4°C.

### Caveolin-1 purification

Washed membranes were resuspended in 4 mL of 50 mM sodium phosphate pH 8.0, 500 mM sodium chloride, 2% (v/v) Empigen BB^®^, and 0.5 mM TCEP + 1 mM dansyl PDS for A87C constructs and 1 mM 1,5 -IAEDANS for F99C constructs. The membrane resuspension was then sonicated using a Branson sonifier 450 equipped with a micro tip (duty cycle 4 output power 5) for 5 min. The reaction occurred at room temperature for 1 hr with steady rotation. The lysate was cleared by centrifugation at 230,000 x g for 20 min at 4°C and filtered through a 0.2 µm PES syringe filter. The clarified lysate was then passed over a column (1 cm diameter) containing 1 mL of Ni -NTA resin equilibrated with 50 mM phosphate pH 8.0, 500 mM sodium chloride, 0.5 % (v/v) Empigen BB^®^. The column was washed with 5 mL of 50 mM phosphate pH 8.0, 500 mM sodium chloride, 0.5% (v/v) Empigen BB^®^, 35 mM imidazole. Cav1 was eluted from the column in 1 mL increments using 50 mM phosphate pH 8.0, 500 mM sodium chloride, 0.5% (v/v) Empigen BB^®^, 250 mM imidazole. Protein-containing fractions (assessed by UV 280) were then combined and concentrated in a Millipore 10kDa MWCO centrifugal concentrator to 1 mL. The sample was filtered through a 0.2 µm PES syringe filter and injected onto a Sephacryl^®^ 300 HR 16/60 column equilibrated with 50 mM phosphate pH 8.0, 150 mM sodium chloride, 0.5% (v/v) Empigen BB^®^, 0.02% sodium azide. Protein purity was assessed using SDS-PAGE. This procedure was repeated aside from dansyl labeling for donor only constructs (W128 and W128 P110A).

### Static Light Scattering

Determination of the average molecular weight can be described using the Debye equation.

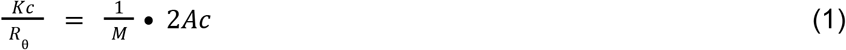

where *K* is an optical constant that is dependent on the refractive index of the solvent (n), the wavelength of light (*λ*_0_), the concentration of protein (c), and the refractive index increment of the protein in solution (dn/dc).

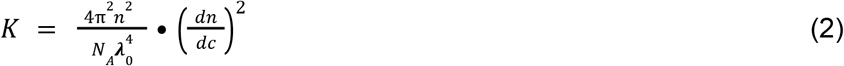

*R*_*θ*_ is the excess Rayleigh scattering ratio. This is directly related to the intensity of the light scattered by the sample solution above that of pure solvent at the given angle, divided by the intensity of the incident beam.

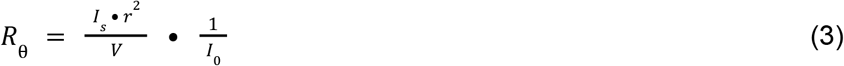

M is the molecular weight of the protein. C is the concentration of the sample, and A_2_ is the second virial coefficient. The second virial coefficient is negligible at low concentrations.

To determine the dn/dc of Cav1 in Empigen BB^®^ micelles, a purified fraction of Cav1 was concentrated in a 50 kDa MWCO Millipore Amicon^®^ Ultra Centrifugal Filter at *100 x g* alongside an equal volume of 50mM sodium phosphate pH 8.0 150mM sodium chloride 0.5% (v/v) Empigen BB^®^ 0.02% sodium azide. Cav1 and the buffer only sample were concentrated to 200 µL. The concentrated samples were quantified at 280 nm with an extinction coefficient of 34,410 M^-1^cm^-1^ (c_protein_ and c_buffer_) The refractive index of each solution (*n*_*protein*_ and *n*_*buffer*_) was measured directly using an Anton Paar Abbemat 3200 benchtop refractometer. dn/dc was calculated using the corrected protein concentration and corrected refractive index, Equation 4. This was repeated in triplicate.

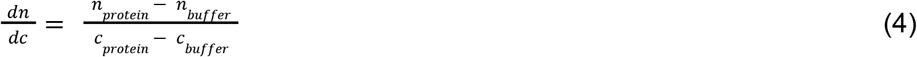

Purified and quantitated Cav1 samples were filtered through a 0.22 µm polyethersulfone (PES) syringe filter, and placed in a small glass test tube (12 mm x 75 mm) for static light scattering analysis. Measurements were performed in quadruplicate for each sample, and each construct was repeated in triplicate. Kc/R is averaged and average molecular weights are summarized in Table 1.

**Table 1.** Summary of weight average molecular mass of Cav1 constructs in Empigen BB^®^ micelles.

| Construct | $Kc/R$ (g/mol)<sup>-1</sup><br>($\times 10^{-5}$) | Weight-Average<br>Molecular Mass (kDa) | Theoretical<br>Molecular Mass<br>(kDa) |
| --- | --- | --- | --- |
| A87C W128 | $4.42 \pm 0.12$ | $22.78 \pm 0.58$ | 22.58 |
| W128 | $4.45 \pm 0.04$ | $22.40 \pm 0.24$ | 22.55 |
| A87C P110A W128 | $4.37 \pm 0.21$ | $22.90 \pm 0.11$ | 22.56 |
| W128 P110A | $4.46 \pm 0.13$ | $22.42 \pm 0.06$ | 22.53 |
| F99C W128 | $4.43 \pm 0.29$ | $22.59 \pm 0.15$ | 22.51 |
| F99C P110A W128 | 4.29 ± 0.56 | 23.32 ± 0.31 | 22.45 |

### Near UV Circular Dichroism Spectroscopy

9µM samples of caveolin-1-myc-H6 C133S C143S C156S with and without the P110A mutation were placed in a 10 mm cuvette and near UV circular dichroism spectra were obtained. A 1 nm bandwidth in step mode accumulating 4 scans from 320 to 260 nm with a 0.5 nm data point interval and 1 second digital integration time. A background spectra of buffer 50 mM sodium phosphate pH 8.0 150 mM sodium chloride and 0.5% empigen BB was subtracted from protein containing spectra. All spectra were collected at 298 K using a JASCO circular dichroism spectrophotometer (Easton, MD) and were repeated in triplicate.

### Determination of Forster Distance for Trp-Dansyl Pair

The Förster distance, *R*_0_ depends on the quantum yield of the donor (*Q*_*D*_), the degree of spectral overlap between the donor emission and acceptor absorption (*J*), the refractive index of the solution (*n*), and the relative orientation in space (κ^2^), following the equation:

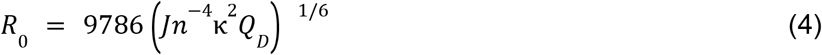

The quantum yield of W128 was determined using a relative method. Quinine sulfate in 0.1 M H_2_SO_4_ was selected as the reference at 25°C. The quantum yield of W128 was determined to be 0.16 ± 0.01. The degree of spectral overlap was calculated as 5.839 × 10^13^ M^-1^ cm^-1^nm^4^. The relative refractive index of the solution was measured as 1.3362 using an Anton Paar Abbemat 3200 benchtop refractometer. Assumed values of ⅔ for κ^2^ were used. The *R*_0_ for the W128 - dansyl pair was calculated as 22.1Å.

### Fraction Labeled of Donor and Acceptor Constructs

Fraction labeled (FL) was determined using Equation 5 (19), where A_337_ and A_280_ represent the absorbance of the sample corrected at 337 nm and 280 nm, respectively. *ε*_*D*337_ represents the extinction coefficient of dansyl at 337 nm which is 5,700 cm^-1^M^-1^. *ε*_*p*280_ is the extinction coefficient of the protein, 18,910 cm^-1^M^-1^.

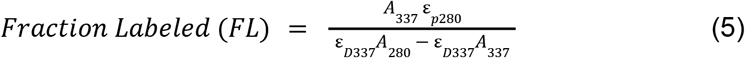

### FRET Efficiency Calculations

FRET efficiency (E) can be calculated using the equation below:

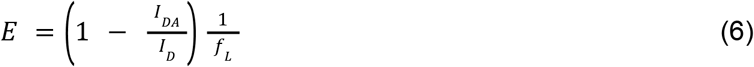

where I_DA_ and I_D_ are the fluorescence intensity in the presence and absence of the acceptor, respectively. f_L_ is the fraction labeled calculated in Equation 5.

### Fluorescence Intensity Measurements

Concentrations of protein pairs were matched using a micro BCA assay. 150 µL of equal concentrations of the donor sample (W128, W128 P110A) and acceptor sample (A87C W128, F99C W128, A87C P110A W128, F99C P110A W128) were placed in a Grenier UV-Star 96 well plate alongside buffer samples. In an Infinite M Nano Tecan plate reader, samples were excited at 295 nm, to avoid tyrosine excitation, and emission was collected at 345 nm, with 100 flashes. Corrected fluorescence intensities were utilized for FRET calculations. All pairs were performed in triplicate alongside W128.

### Monitoring FRET from 5ºC to 45ºC

Emission spectra from 305 to 450 nm with an excitation at 295 nm were collected for A87C W128 and F99C W128 donor and donor-acceptor constructs over 5 temperatures. The temperature was varied by increments of 10*º*C from 5*º*C to 45*º*C using a Horiba Fluorolog FL3-22iHR spectrophotometer with a temperature-controlled module. The temperature was allowed to equilibrate for 5 minutes at each temperature before spectra were obtained. All spectra were collected using a cuvette with a 1 cm path length.

## Results and Discussion

To investigate the intramembrane turn, the tertiary structure of wild-type Cav1 was compared to that of the P110A mutant using near-UV circular dichroism (CD) spectroscopy. Near-UV CD is a technique that reports on the chiral environment of the aromatic side chains, providing information about the tertiary structure of the protein. CD signals in the near-UV range (320-260 nm) arise from the orientation of aromatic residues within a folded protein and the intensity is sensitive to changes in the tertiary structure. Here, near-UV CD revealed a dramatic reduction in molecular ellipticity at 280 nm in the P110A mutant compared to wild-type Cav1, suggesting changes in the tertiary structure (Figure 3). 280 nm primarily reflects the chiral environment of aromatic residues, particularly tryptophan and tyrosine, which are spread relatively evenly across the critical helices 1 and 2. The observed loss of ellipticity indicates that the P110A mutation likely disrupts the tertiary arrangement of helices 1 and 2. Futhermore increased sidechain flexibility is consistent with linearizing the helix (Figure 2A) as opposed to maintaining the U-shaped topology and simply flipping Cav1’s orientation in the membrane.

**Figure 2.**
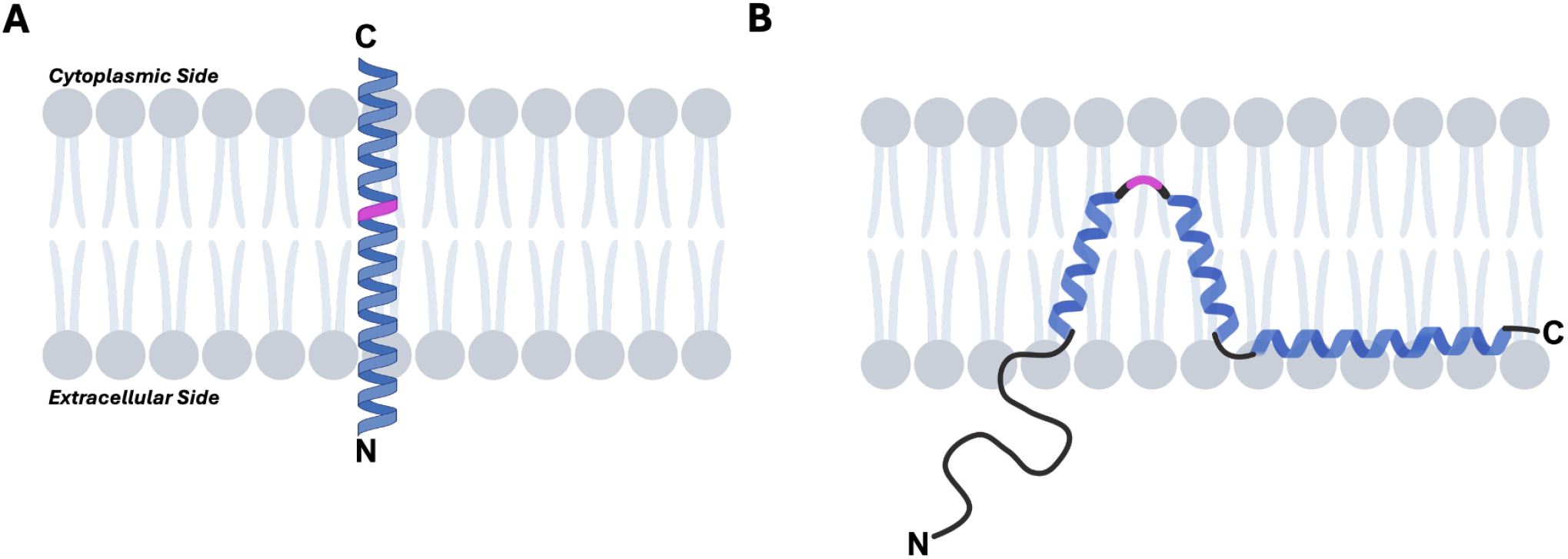
***A***. Hypothesized linearization of Cav1 resulting in extracellular N-terminus. ***B.*** Alternative hypothesized topology of Cav1 where U-shaped conformation flips in the membrane. P110 is highlighted in pink.

**Figure 3.**
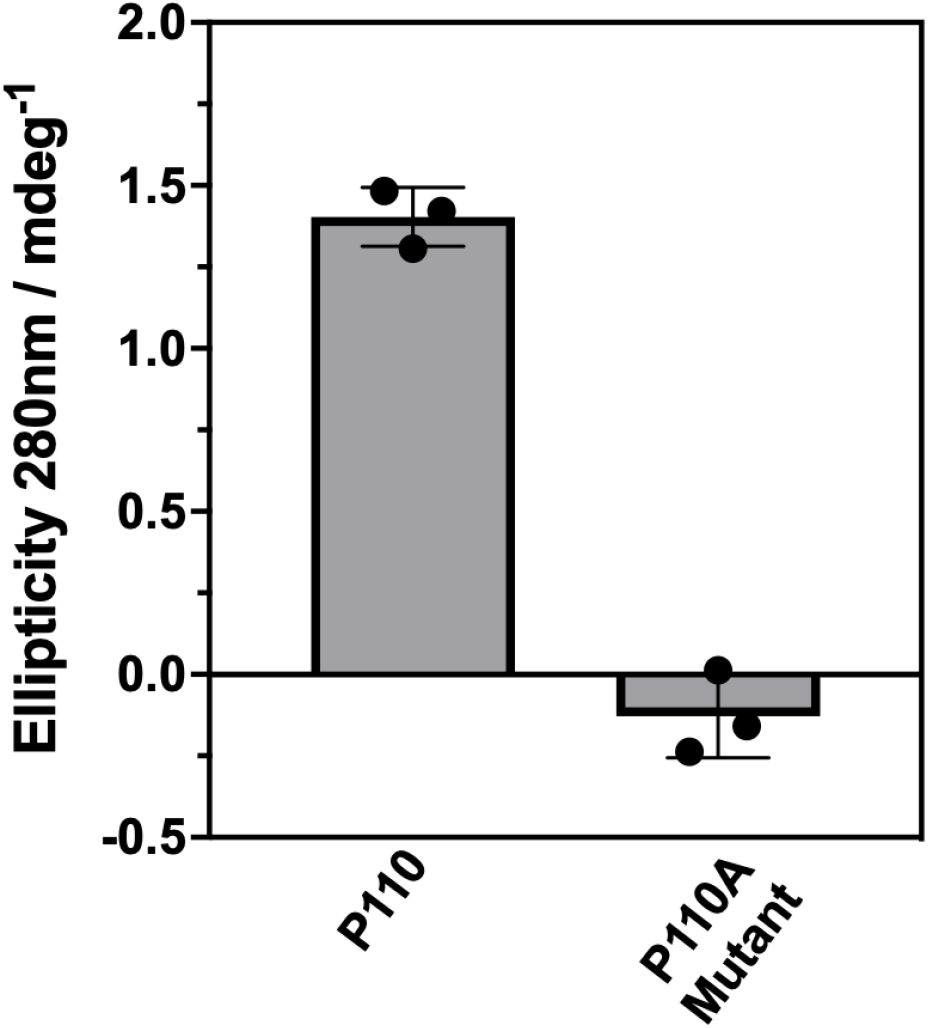
N*ear-UV circular dichroism analysis of wild-type Cav1 (P110) and the P110A mutant.* Molecular ellipticity at 280 nm was measured by near-UV circular dichroism spectroscopy. Bars represent mean ellipticity values ± SEM (n = 3).

The tryptophan-dansyl pair was determined to have an R_0_ of 22.1Å, allowing FRET to be observed from 11Å - 44Å. The donor position, W128, rests at the C-terminus of H2 while the acceptor position, A87C or F99C, rests at the N-terminus or center of H1, respectively (Figure 1). These positions allow for a way to directly monitor drastic changes in FRET efficiencies if conformational changes occur. In the event that the P110A mutation induces linearization of H1 and H2 into a single continuous transmembrane helix, the spatial separation between donor and acceptor residues would increase substantially, resulting in a pronounced decrease in FRET efficiency. Specifically, assuming a fully linearized helix conformation, the distance between residues 87 and 128 is estimated to be ∼58.5 Å, exceeding twice the Förster radius (2R_0_) and thus precluding detectable FRET. To probe shorter distances within the FRET distance range upon potential linearization, an alternative acceptor site was introduced at residue 99 (F99C), which would reduce the maximum donor-acceptor distance to approximately 43.5 Å, enabling measurement of low-efficiency FRET states.

An important consideration in our FRET studies was to ensure that the protein was monomeric. This avoided any confusion as to whether the FRET was arising from unwanted intermolecular FRET. Static light scattering (SLS), is an excellent technique to quantitatively determine the weight-averaged molecular mass of a protein sample (21). Briefly, the molecular weight of a molecule is directly proportional to the magnitude of scattered light. Obtaining Kc/R values requires calculating the dn/dc; i.e., the change in refractive index per change in protein concentration, of the protein-detergent complex. The dn/dc of Cav1 in Empigen BB^®^ was determined to be 0.259 ± 0.002 mL/g. Table 1 shows the determined molecular weights of the two FRET pairs along with the donor-only, W128 and W128 P110A, constructs.

The weight-averaged molecular mass for each construct is within 3% of the actual molecular weight of Cav1, showing that Cav1 in Empigen BB^®^ micelles is highly monomeric. The advantage of SLS as a method for the determination of membrane protein oligomeric states is that it is highly sensitive, any significant amount of aggregation would have drastically reduced the Kc/R value and hence increased the weight-averaged molecular weight.

### Observed FRET Efficiencies forThe FRET pairs P110 and P110A mutant

Experimentally, the FRET efficiency between A87C and W128 was determined to be **56.2 ± 0.4 %** (Figure 4A, P110). Given that the acceptor fluorophore is positioned at the N-terminal end of H1 (A87C), and the donor is positioned at the C-terminal end of H2 (W128), 42 amino acids separate them. Based on previous NMR studies from our lab, of those 42 residues only three, **GIP**, are excluded from ***α***-helical structure (14,22). Therefore, if H1 and H2 are ideal ***α***-helices extended linearly, the efficiency between the donor and the acceptor fluorophore would not be within the limit of detection. The A87C W128 mutant results in 56% FRET efficiency, which would require Cav1 to adopt an intramembrane turn bringing the residues into closer proximity. When proline at position 110 is mutated to alanine (P110A) in A87C W128, FRET is nearly eliminated, dropping to **0.44 ± 0.04 %**, which falls beyond the 2R_0_ detection limit. This significant loss in efficiency strongly supports the hypothesis that the P110A mutation induces a major conformational change, via linearization of helices H1 and H2.

**Figure 4.**
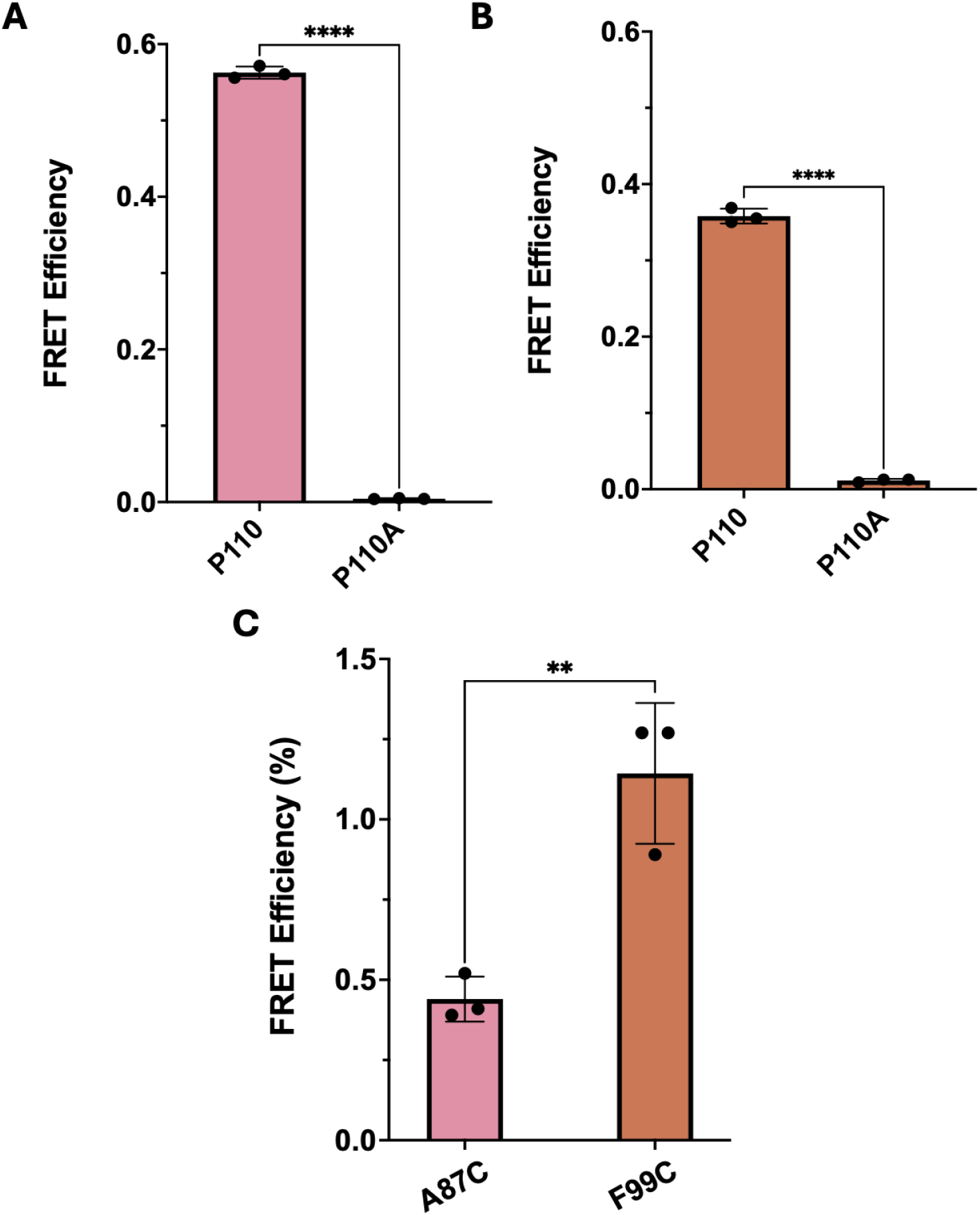
***A.*** FRET efficiency measurements for the A87C W128 construct in the presence of P110 and P110A. ***B.*** FRET efficiency measurement for F99C W128 under the same conditions. Data shown as mean ± SEM (n = 3). ***C.*** Comparison of percent FRET efficiency between A87C P110A W128 and F99C P110A W128.

### Observed FRET Efficiencies for F99C W128 WT and P110A

Given that the P110A mutation in A87C W128 results in a significant loss of FRET that it is beyond the 2R_0_ detection limit, we designed a second construct to restore measurable FRET efficiency in the presence of the P110A mutation. To achieve this, we repositioned the acceptor fluorophore further along helix 1 at residue 99 by mutating phenylalanine 99 to cysteine (F99C), while retaining W128 as the donor position. *Figure 4B* shows a FRET efficiency of **35.8 ± 0.6 %**, representing a significant decrease compared to the A87C W128 pair, consistent with an increased distance between fluorophores expected if the protein adopts an intramembrane turn. Upon introduction of the P110A mutation, FRET efficiency further decreases significantly to **1.14 ± 0.1%**. Together, these results suggest that the P110A mutation induces a major conformational change in Cav1, extending the helices as evidenced by the dramatic loss of FRET efficiency. Importantly, the more modest decrease in FRET efficiency observed in the F99C W128 construct compared to the A87C W128 pair upon P110A mutation is consistent with a model where the helices are linearized rather than forming a compact intramembrane turn.

### Assessment of molecular motion: monitoring FRET distance across temperatures

Molecular motion can impact FRET by altering the distance and orientation of the fluorophores. To evaluate if the molecular motion had any consequential impacts on the efficiencies observed, the FRET efficiency between both A87C W128 and F99C W128 was monitored from 5ºC to 45ºC. As temperature rises, molecular motion increases; thus, significant changes in FRET efficiency with rising temperature would suggest that the measured FRET efficiencies at ambient temperature may not accurately reflect the native conformation of Cav1. However, *Table 2* demonstrates consistent FRET efficiencies across a large temperature range, indicating that molecular motion does not substantially alter the conformation of Cav1. The intensity of the tryptophan fluorescence decreased with temperature, likely due to enhanced quenching from more frequent collisions between tryptophan residues and solvent molecules.

**Table 2.** Temperature-dependent fluorescence intensities and FRET efficiencies for the A87C W128 and F99C W128 Cav1 constructs. Donor fluorescence intensity was measured in the absence and presence of the acceptor fluorophore across temperatures ranging from 5ºC to 45ºC. FRET efficiency was calculated at each temperature to assess the impact of increased molecular motion on energy transfer and protein conformation.

| ***A87C W128*** | **5°C** | **15°C** | **25°C** | **35°C** | **45°C** |
| --- | --- | --- | --- | --- | --- |
| **Fluorescence Intensity - Donor** | 1.44 x 10<sup>6</sup> | 1.40 x 10<sup>6</sup> | 1.29 x 10<sup>6</sup> | 1.12 x 10<sup>6</sup> | 9.78 x 10<sup>5</sup> |
| **Fluorescence Intensity - Donor + Acceptor** | 8.60 x 10<sup>5</sup> | 8.05 x 10<sup>5</sup> | 7.57 x 10<sup>5</sup> | 6.55 x 10<sup>5</sup> | 5.67 x 10<sup>5</sup> |
| **FRET Efficiency** | 0.569 | 0.600 | 0.584 | 0.587 | 0.594 |
| ***F99C W128*** | **5°C** | **15°C** | **25°C** | **35°C** | **45°C** |
| **Fluorescence Intensity - Donor** | 8.16 x 10<sup>5</sup> | 7.39 x 10<sup>5</sup> | 6.61 x 10<sup>5</sup> | 6.17 x 10<sup>5</sup> | 5.12 x 10<sup>5</sup> |
| **Fluorescence Intensity - Donor + Acceptor** | 5.90 x 10<sup>5</sup> | 5.44 x 10<sup>5</sup> | 4.93 x 10<sup>5</sup> | 4.36 x 10<sup>5</sup> | 3.76 x 10<sup>5</sup> |
| **FRET Efficiency** | 0.366 | 0.349 | 0.336 | 0.388 | 0.352 |

## Conclusion

Computational experiments previously suggested the mutation of proline 110 to alanine (P110A) could drastically alter the Cav1 topology by converting a native transmembrane turn into a linear transmembrane helix. However, this remained unclear experimentally and raised the question of whether P110A causes alpha helical linearization or another distinct topological rearrangement that results in an extracellular N-terminus. To address this ambiguity and provide experimental insight into the helical arrangement of Cav1, we employed FRET between a native tryptophan and a site-specific dansyl probe. The observed FRET data reveals that mutation of P110A causes a major conformation shift that alters the spatial relationship between H1 and H2, resulting in a dramatic decrease in FRET efficiency. This supports a model where proline 110 is essential for maintaining a native intramembrane turn of Cav1, and upon mutation results in an extended transmembrane helix. These results provide new experimental support for the hypothesized V-shaped arrangement of Cav1 and highlight the structural importance of P110, further advancing our understanding of caveolin topology.

## Acknowledgements

Figure 1 and Figure 2 were created using BioRender.com. This work was supported by the NIH R15 GM141606-01 awarded to Kerney Jebrell Glover and Wonpil Im. The authors thank Drs. Christophe Guillon and Jaya Saxena for the use of their Infinite M Nano Tecan plate reader.

